# Assembly and early maturation of large subunit precursors

**DOI:** 10.1101/486225

**Authors:** Malik Chaker-Margot, Sebastian Klinge

**Affiliations:** Laboratory of Protein and Nucleic Acid Chemistry, The Rockefeller University, New York, NY, US; Tri-Institutional Training Program in Chemical Biology. The Rockefeller University, New York, NY, US

## Abstract

The eukaryotic ribosome is assembled through a complex process involving more than 200 factors. As pre-ribosomal RNA is transcribed, assembly factors bind the nascent pre-rRNA and guide its correct folding, modification and cleavage. While these early events in the assembly of the small ribosomal subunit have been relatively well-characterized, assembly of the large subunit precursors, or pre-60S, is less well understood. Recent structures of nucleolar intermediates of large subunit assembly have shed light on the role of many early large subunit assembly factors but how these particles emerge is still unknown. Here, we use the overexpression and purification of truncated pre-rRNAs to examine the initial assembly of pre-60S particles. Using this approach, we can recapitulate the early recruitment of large subunit assembly factors mainly to the domains I, II and VI of the assembling 25S rRNA.

## Introduction

Ribosome assembly in eukaryotes is a complex and intricate process which requires more than 200 non-ribosomal factors in yeast (Woolford and Baserga 2013). Ribosome biogenesis begins in the nucleolus where RNA polymerase I transcribes the 35S pre-rRNA, a polycistronic RNA, which contains the 18S, 5.8S and 25S rRNAs. These rRNAs are flanked by 5’ and 3’ external transcribed spacers (ETS) and two internal transcribed spacers (ITS), ITS1 and ITS2. The 5S rRNA is transcribed separately by RNA polymerase III as a pre-5S RNA species. As RNA polymerase I transcribes the 35S pre-rRNA, ribosome assembly factors bind the nascent pre-rRNAs forming large multi-component complexes, which represent the earliest precursors of the small and large ribosomal subunit. Cleavage of the pre-rRNA at the sites A2 and A3, located within ITS1, separate small and large subunit pre-rRNA and thereby small and large subunit assembly.

Recent structures of nucleolar pre-60S particles have revealed some of the folding pathways that large subunit pre-rRNA follows (Kater et al. 2017; Sanghai et al. 2018; Zhou et al. 2018) showing that domains I, II and VI of the 25S rRNA, along with the 5.8S rRNA, fold first into a near-mature conformation. Domains III-V remain unfolded in the early stages of large subunit assembly.

Despite the wealth of information provided by these new structures, it remains unclear how nucleolar pre-60S particles emerge. The recruitment of ribosome assembly factors on pre-rRNA has been elucidated for the first part of the pre-rRNA using the overexpression of truncated pre-rRNA to recapitulate this process (Chaker-Margot et al. 2015; Zhang et al. 2016). This has led to the description of the assembly of the earliest precursor of the small subunit. More recently, another group has used the same approach to recapitulate the assembly of large subunit precursors onto pre-rRNA (Chen et al. 2017). Here, using a similar technique, we described our model for early large subunit assembly.

## Results

We set out to determine the order in which large ribosomal subunit assembly factors are recruited to the second half of the nascent pre-rRNA. For this purpose, purification of pre-ribosomal complexes relies on the overexpression of tagged truncated pre-rRNAs, which are then bound by endogenous ribosome assembly factors, and their isolated in a two-step affinity purification, as previously (Chaker-Margot et al. 2015). We co-expressed the truncated pre-rRNAs with 5 repeats of the MS2 aptamers at the 3’ end together with a protease-cleavable MS2-GFP fusion (Fig. 1A). The MS2 protein is fused to a nuclear localization signal (NLS) to promote its association with the truncated RNA in the cell. The expressed and truncated pre-rRNAs are recognized by the endogenous ribosome assembly machinery and isolated in a two-step purification (Fig 1.B).

**Figure 1.**
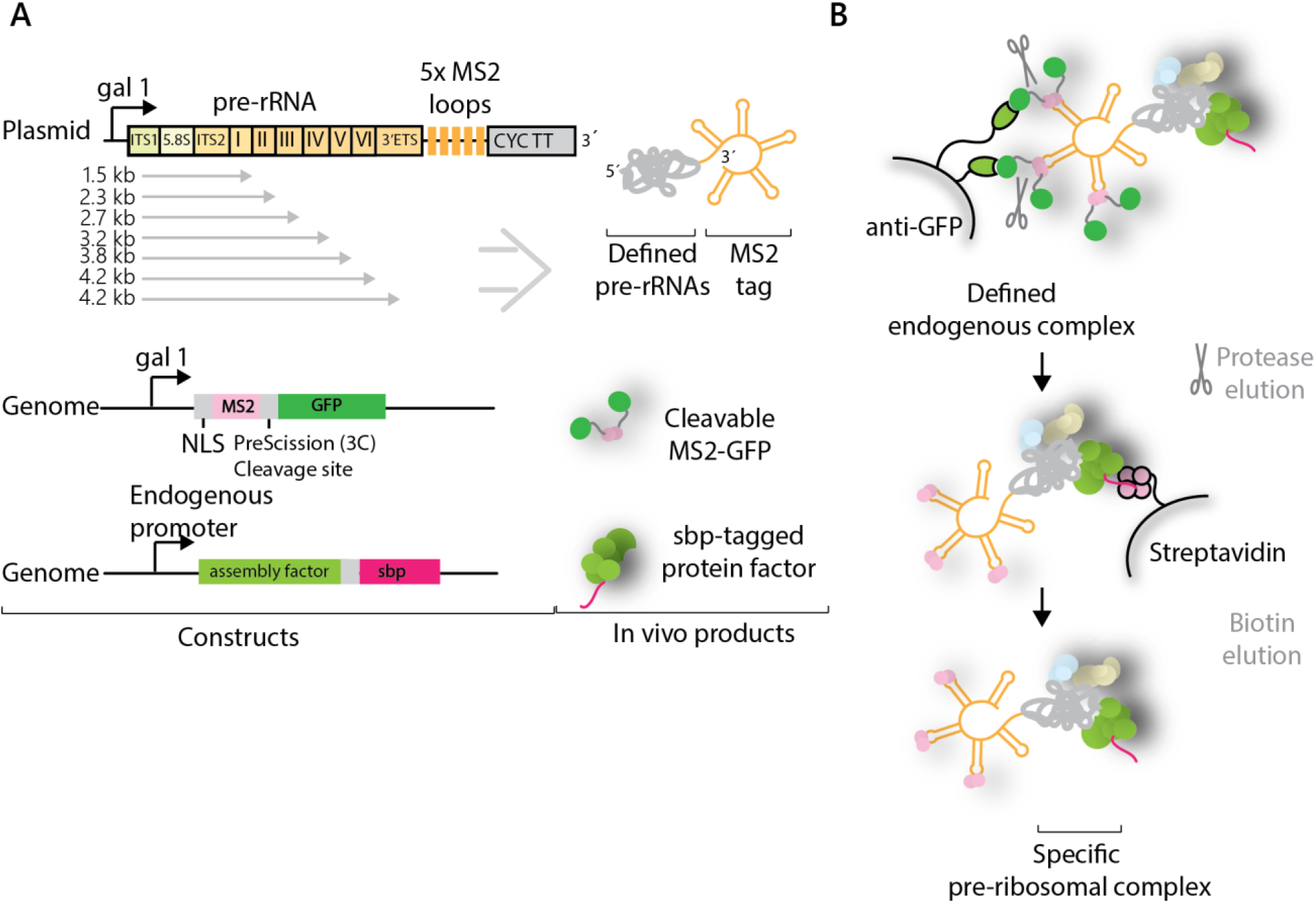
Purification of pre-ribosomal particles. (**A**) Plasmid overexpression of aptamer-tagged pre-rRNAs of different lengths in genetically modified yeast strains containing a sbp-tagged ribosome assembly factor. Plasmids contain the gal1 promoter and CYC transcriptional terminator (CYC TT). Pre-rRNA domains, sbp-tagged ribosome assembly factor and MS2-GFP protein are shown. (**B**) Purification strategy using GFP and sbp as sequential tags, followed by biotin elution.

The RNA constructs used here begin at the A3 site thereby mimicking A3 cleaved pre-rRNAs. Previous studies have suggested that separation of the rDNA locus between the A2 and A3 sites, at nucleotides 2739 to 2747 of the 35S pre-rRNA, leads to viable ribosome production suggesting that there is no essential link between small and large subunit assembly factors (Liang and Fournier 1997). Target RNAs span the second half of the 35S locus, ending at each of the structural domains of the 25S, I-VI (Fig. 1A). The final and longest construct spans the entirety of the 25S and ends at the B0 site. A “full-length” construct that ended at the native end of the pre-rRNA underwent B0 cleavage, thereby separating the 3’ MS2 tag from the rest of the RNA and making isolation impossible (data not shown). The ITS2-binding nucleolar factor Cic1 was chosen as protein bait for the second capture (Wu et al. 2016). By using Cic1 as protein bait, we ensured that ITS2 processing has not yet occurred, thereby narrowing the type of pre-60S particles that can be isolated to earlier species.

Upon induction, truncated pre-rRNAs are expressed and bound by MS2-GFP and the endogenous large subunit assembly machinery, including the sbp-tagged Cic1. During the purification, the RNA molecules are captured by the anti-GFP resin (Fig. 1B). In the second step, the targeted pre-ribosomal complex is enriched by streptavidin capture, and subsequent biotin elution.

The overexpressed truncated pre-rRNAs were analyzed in total RNA extracts by northern blotting using an MS2 probe (Suppl. Fig. 1A). RNA bands corresponding to their expected size were visualized for each sample but were weak for the domain VI and B0 truncated species. Surprisingly, the major bands for the domain VI and B0 species run lower that the domain V band. These bands could represent degradation or processing products of these species and have been observed previously (Chen et al. 2017). Since the MS2 loops are located at the 3’ end of the constructs, this allows mapping the approximate cleavage site to around nucleotide 1350 in domain II. The mechanism by which this cleavage happens is unknown.

The use of our purification system has allowed us to recapitulate the assembly of the earliest large subunit assembly intermediates. The negative control, wherein only the MS2 loops were expressed, yielded an accordingly clean sample, except for the presence of streptavidin (Fig. 2A). The addition of the first section of pre-rRNA, spanning from the A3 site to the end of domain I of the 25S, lead to the recruitment of approximately 10 factors. The subsequent addition of domains of the 25S induced the recruitment of more factors, although in a less dramatic fashion than what was observed for the small subunit processome (Chaker-Margot et al. 2015). Samples for all seven constructs, with termini at domain I-VI and B0, plus a negative control were purified in biological triplicates and analyzed by LC-MS. The median amounts (as quantified by areas) for each protein were then used for the data analysis (Fig. 2B, C). Only proteins with amounts within 100-fold of the protein bait Cic1, and with more than 5 unique peptides were considered for the analysis. The amounts of Cic1, which should stay similar throughout, were used to normalize the samples. As previously (Chaker-Margot et al. 2015), ribosomal proteins have been excluded from this analysis, as signals of highly abundant peptides from contaminating mature ribosomes prevent an unambiguous assignment of ribosomal proteins to a particular stage of ribosome assembly in our system.

**Figure 2.**
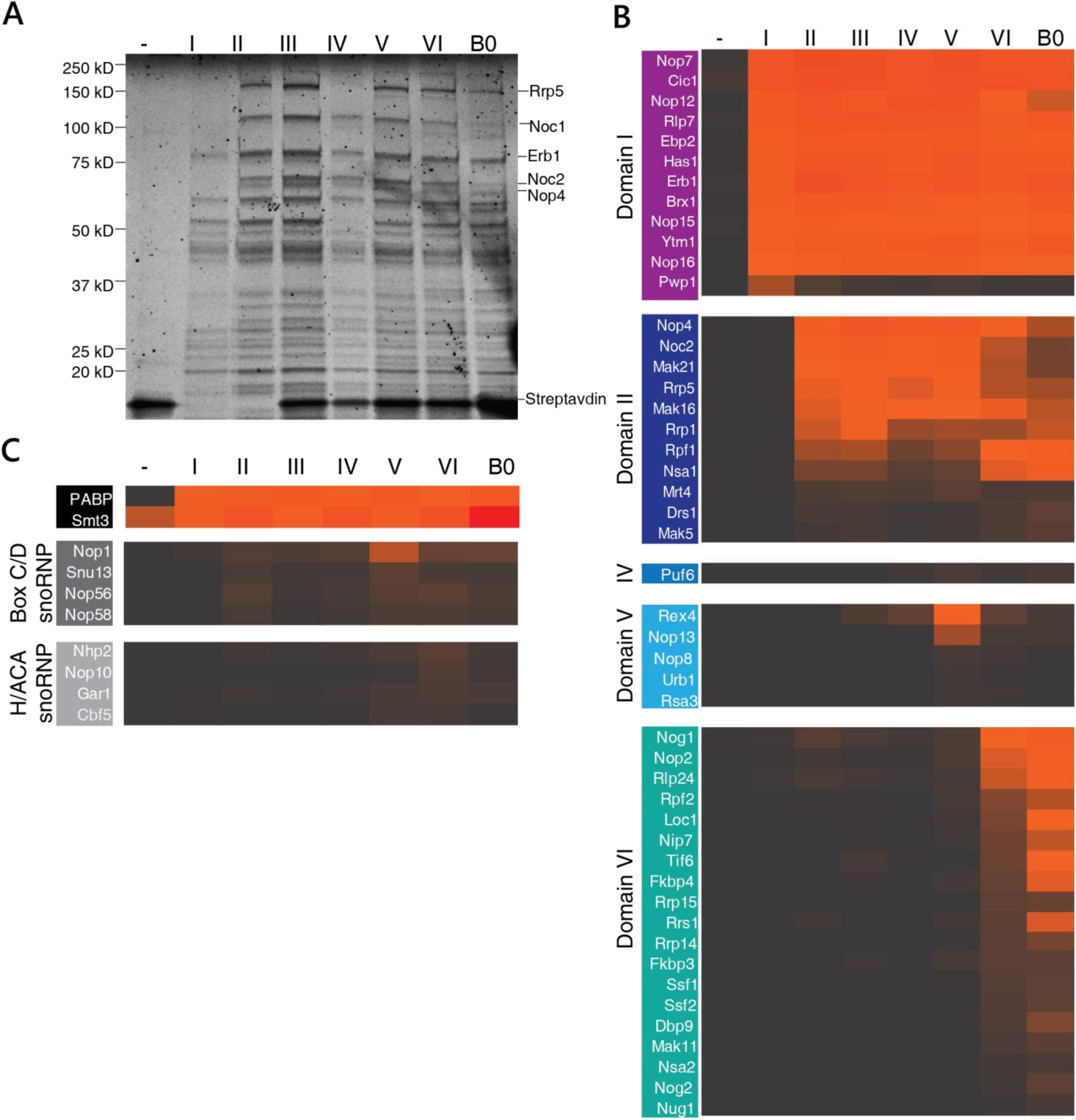
Stage-specific assembly of nucleolar pre-60S particles. (**A**) SYPRO-Ruby stained SDS-PAGE gel of pre-ribosomal complexes corresponding to increasing length of large subunit pre-RNA. Ribosome assembly factors are indicated on the right. (**B**) LC-MS analysis of ribosome assembly factors, based on a triplicate of samples, normalized against Cic1 and classified by recruitment stage. Protein abundance is measured by peak area and high abundance proteins are shown in orange/red while low abundance and absent proteins are shown in black/grey. (C) LC-MS analysis of the polyadenylate binding protein (PABP), Smt3 and the H/ACA and box C/D snoRNP proteins.

This analysis has allowed us to determine the timing of recruitment of approximately 50 assembly factors on nascent pre-rRNA (Fig. 2B, C). The first construct (containing 5.8S rRNA, ITS2 and domain I) recruits 12 assembly factors, including several known binders of ITS2 (Cic1, Nop7, Nop15 and Rlp7) (Wu et al. 2016). The helicase Has1, which is involved in small and large subunit processing (Emery et al. 2004) is also incorporated in this particle. Strikingly, this first construct also induces the recruitment of the protein Pwp1, whose levels decrease drastically for longer constructs, indicating its transient role in early large subunit assembly (Talkish et al. 2014). High levels of the ubiquitin like modifier SUMO (Smt3) were detected in all intermediates, suggesting a possible functional role in early large subunit assembly (Fig. 2C) (Panse et al. 2006; Raman et al. 2016).

Extension of the pre-rRNA into domain II recruits another 11 factors. Importantly, and in contrast to previous data (Chen et al. 2017), these include the large protein Rrp5 and its known interaction partners, Noc1 and Noc2. Rrp5 is required for the assembly of the small subunit while Noc1/Noc2 have been shown to co-precipitate with the SSU processome (Chaker-Margot et al. 2015), likely through their interaction with Rrp5 (Hierlmeier et al. 2013). *In vivo* crosslinking studies of Rrp5 have shown its proximity to several sites on the pre-rRNA, including the central domain of the 18S, ITS1 and domain II of the 25S, consistent with our data (Lebaron et al. 2013). Domain II also leads to the association of several factors completing the ring structure around domain I and II which was described in the nucleolar pre-60S structure (Sanghai et al. 2018), composed of Nsa1, Rpf1, Mak16, Rrp1, and the already associated Brx1 and Ebp2.

While the addition of domain III recruits no specific assembly factors, domain IV induces the binding a single protein, Puf6. Domain V leads to the appearance of Nop13, a known nucleolar pre-60S factor and 3 members of the Npa/Urb subcomplex: Urb1, Nop8 and Rsa3. These proteins form a subcomplex, which also includes Urb2 and the helicase Dbp6, involved in the earliest steps of large subunit biogenesis (Rosado et al. 2007; Joret et al. 2018). While its role in large subunit assembly is still poorly understood, this data suggests that this complex associates with domain V.

Completion of the 25S rRNa induced the recruitment of many factors, including the GTPase Nog1, which remains associated with the pre-60S particle until its removal in the nucleus, as well as Tif6 and Rlp24, which are removed in the cytoplasm. Several nucleolar specific factors also bound at this stage, such as Nop2, Rrp14, the Ssf1-Rrp15 dimer and Nip7. Surprisingly, low levels of Nog2 were also detected in the A3-domain VI sample (Fig. 2B). Nog2 is a nuclear assembly factor whose binding site in the pre-60S particles overlaps with nucleolar factors such as Nip7, Sbp1 and Nop2 (Wu et al. 2016; Kater et al. 2017), which makes its association with nucleolar intermediates unlikely. This may be evidence that the A3-domain VI sample represents multiple pre-60S intermediates, which have matured to different extents rather than a specific large subunit precursor. Importantly, we did not detect the presence of Nop53, which is now known to have overlapping binding sites with Erb1 and is required for ITS2 processing at later stages of maturation (Thoms et al. 2015; Kater et al. 2017; Sanghai et al. 2018).

Maturation of large subunit precursors involves the recruitment of the 5S rRNA, as well as the modification of its pre-rRNA guided by small nucleolar RNAs. We therefore tested the presence of such RNAs in each of the samples. Northern Blotting analysis of the samples showed that the 5S rRNA appears with the addition of domain VI (Suppl. Fig. 1B). Consistent with this result, we observed that the chaperones of the 5S rRNA, Rpf2 and Rrs1, are also recruited at domain VI.

snR10, which is involved in the pseudouridylation of U2923, in domain V near domain VI (Ni et al. 1997), is also recruited to domain VI (Suppl. Fig. 1B). Correspondingly, there was a moderate increase in signal in the levels of box H/ACA proteins (Cbf5, Nhp2, Nop10 and Gar1) at domain VI, consistent with the presence of snR10. The presence of both 5S rRNA and snR10 is qualitative only, since the truncated pre-rRNAs could not be used as loading controls due to degradation (data not shown).

By overexpressing truncated pre-rRNAs, we have recapitulated the assembly of ribosome biogenesis factors on the second half of the pre-rRNA locus. Large subunit assembly begins with the recruitment of many factors, which bind ITS2, 5.8S and domain I. With the addition of domain II, the Rrp5-Noc1-Noc2 complex, which is also associated with the small subunit processome, binds the forming large subunit, thereby bridging small and large subunit assembly (Fig. 3). Further transcription does not lead to addition of many additional factors, since domains III-V are flexible in the early stages of pre-60S maturation (Suppl. Fig. 2) (Kater et al. 2017; Sanghai et al. 2018; Zhou et al. 2018). Completion of domain VI induces the incorporation of almost 20 ribosome assembly factors, the 5S rRNA and association of snR10. Many of these factors will stay associated with the pre-60S particle until their removal in the nucleus or the cytoplasm. In contrast, the Rrp5-Noc1-Noc2 complex dissociates from these early precursors before their exit from the nucleolus (Fig. 3).

**Figure 3.**
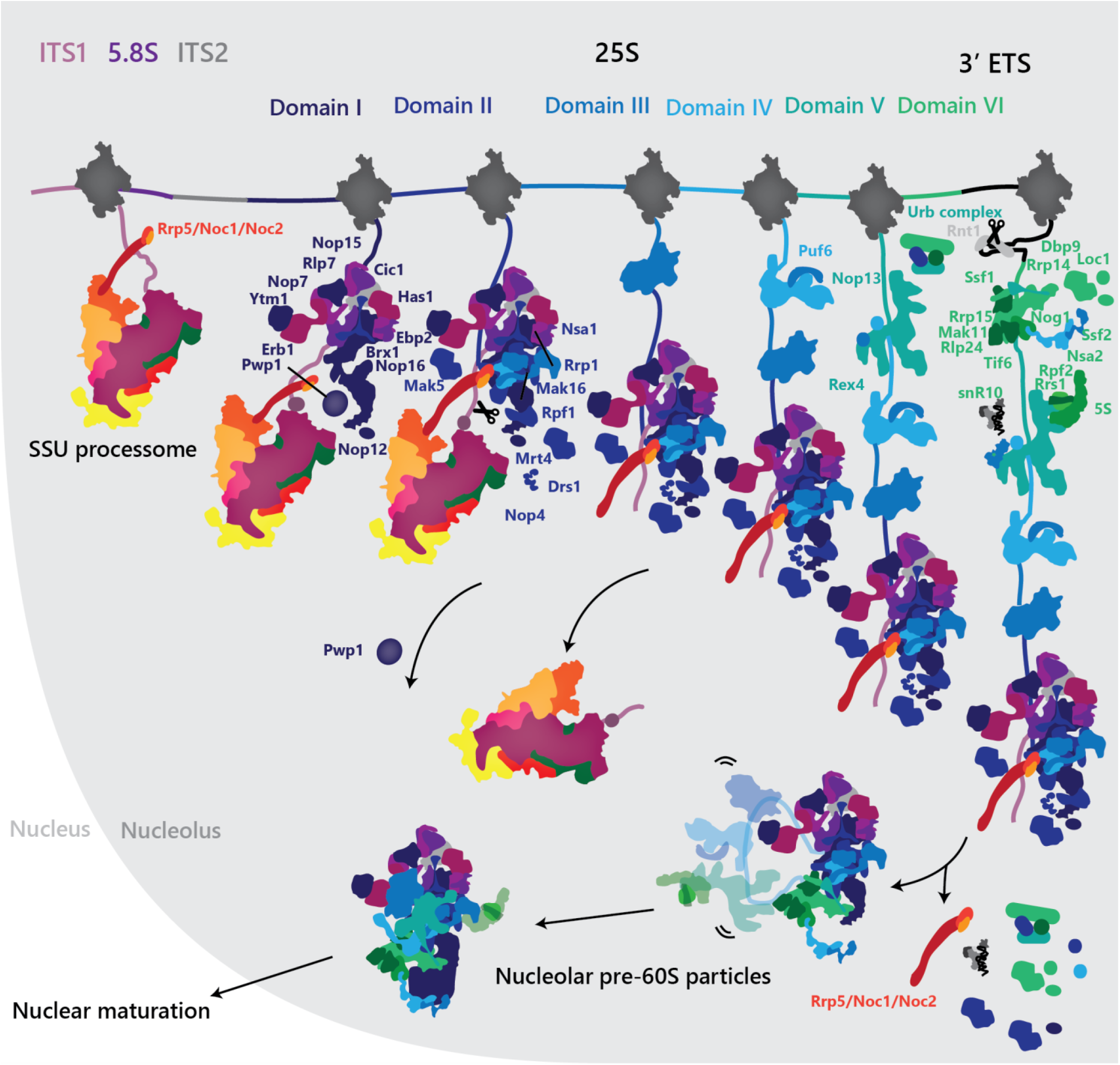
Assembly of the nucleolar pre-60S particle. Updated model of the early assembly of large ribosomal subunit precursors. Step-wise recruitment of ribosome assembly factors is concluded by cleavage of the B0 site by Rnt1 and release of the emerging pre-60S. Re-arrangement of this particle induces the departure of several factors and the formation of the nucleolar pre-60S that was observed previously.

Maturation of large subunit precursors requires processing at the 5’ and 3’ of the precursor RNA, in ITS1 and the 3’ ETS. This processing involves cleavage and resection of the spacer RNAs by Rnt1 and Rex1 for the 3’ end and Rrp17 or Rat1 for the 5’ end (Henry et al. 1994; van Hoof 2000; Oeffinger et al. 2009). It is tempting to speculate that the removal of Rrp5-Noc1-Noc2 may be coupled to 5’ and 3’ processing of the pre-rRNA. Ongoing and future studies will elucidate the precise mechanisms by which these early precursors mature to form nucleolar pre-60S particles.

## Methods

### Cloning of RNA sequences

Ribosomal RNA sequences were amplified directly for BY4741 yeast genomic DNA. Sequences corresponding to all constructs used here from *Saccharomyces cerevisiae* rDNA were cloned downstream of the Gal1 promoter of a derivative of pESC URA (Agilent Technologies) containing five copies of the MS2 aptamer tag at the 5’ or 3’ end.

### Purification of pre-ribosomal complexes assembled on truncated pre-rRNA

Large ribosomal subunit precursors were purified from yeast strains bearing one sbp tag on Cic1 transformed with the appropriate plasmid. The yeast was grown in fully synthetic medium supplement with 2% raffinose and induced with 2% galactose 16hrs. The cells were then harvested and cryogenically ground (Retsch PM100 CM). Cryo-ground yeast powder from was resuspended in buffer A (50 mM Tris-HCl, pH 7.7 (20 °C), 150 mM NaCl, 1 mM EDTA, 0.1% Triton-X100, PMSF, Pepstatin, E-64), and the lysate cleared by high-speed centrifugation. The cleared lysate was incubated with anti-GFP beads. After washes in buffer A, the immobilized complexes were incubated with TEV protease and the supernatant was applied to streptavidin beads. Beads were subsequently washed in buffer B (50 mM Tris-HCl pH 7.7 (20 °C), 150 mM NaCl, 1 mM EDTA) and the complex was eluted in the same buffer, supplemented with 5 mM D-Biotin. Composition of the particles processome was analyzed on a 4-12% SDS-PAGE gradient gel and by LC-MS.

### Northern Blotting

RNA was extracted from either approximately 0.1 g of yeast powder, or 200 uL of biotin eluates, which were resuspended in 1 mL of TRIzol (Life Technologies) and processed according to the manufacturer’s instructions. Total RNA samples are separated on a denaturing 3.7% Formaldehyde - 1.2% Agarose gel (SeaKem LE, Lonza). The RNA was subsequently transferred onto a cationized nylon membrane (Zeta-Probe GT, Bio-Rad) using downward capillary transfer. RNA was cross-linked to the membrane for northern blot analysis by UV irradiation with a UV Stratalinker 2400 (Stratagene). Cross-linked membranes were incubated with hybridization buffer (750 mM NaCl, 75 mM trisodium citrate, 1 % (w/v) SDS, 10% (w/v) dextran sulfate, 25% (v/v) formamide) at 65 °C for 30 min prior to addition of γ-^32^P-end-labeled DNA oligo nucleotide probes.

To generate γ-^32^P-end-labeled DNA oligo nucleotide probes, 1.5-2 μL of DNA oligonucleotide was incubated with 4 μL of γ-32P-ATP and 1 μL T4 PNK in 20 μL total T4 PNK buffer (NEB). The reaction was incubated at 37 °C for 30 min and subsequently cleaned-up by using the illustra MicroSpin G-25 columns (GE). Probes were allowed to hybridize at 65 °C for 1 hour and then at 37 °C overnight. Membranes were washed once with wash buffer 1 (300 mM NaCl, 30 mM trisodium citrate, 1% (w/v) SDS) and once with wash buffer 2 (30 mM NaCl, 3 mM trisodium citrate, 1% (w/v) SDS) for 20 min each at 45 °C. Radioactive signal was detected by exposure of the washed membranes to a storage phosphor screen which was scanned with a Typhoon 9400 variable-mode imager (GE Healthcare).

## Acknowledgements

We thank Jonas Baradun, Mirjam Hunziker, Zahra Sanghai and Linamarie Miller for critical advice in designing the experiments. We also thank Henrik Molina, Milica Tesic Mark and Brian D. Dill from the Rockefeller proteomics core facility for the processing of mass spectrometry samples. M.C-M. was supported by an NSERC PGS D fellowship. S.K. is supported by the Robertson Foundation, the Irma T. Hirschl Trust, the Alexandrine and Alexander L. Sinsheimer Fund, the Rita Allen Foundation, and an NIH New Innovator Award (1DP2GM123459).

---

**Supplemental Figure 1.**
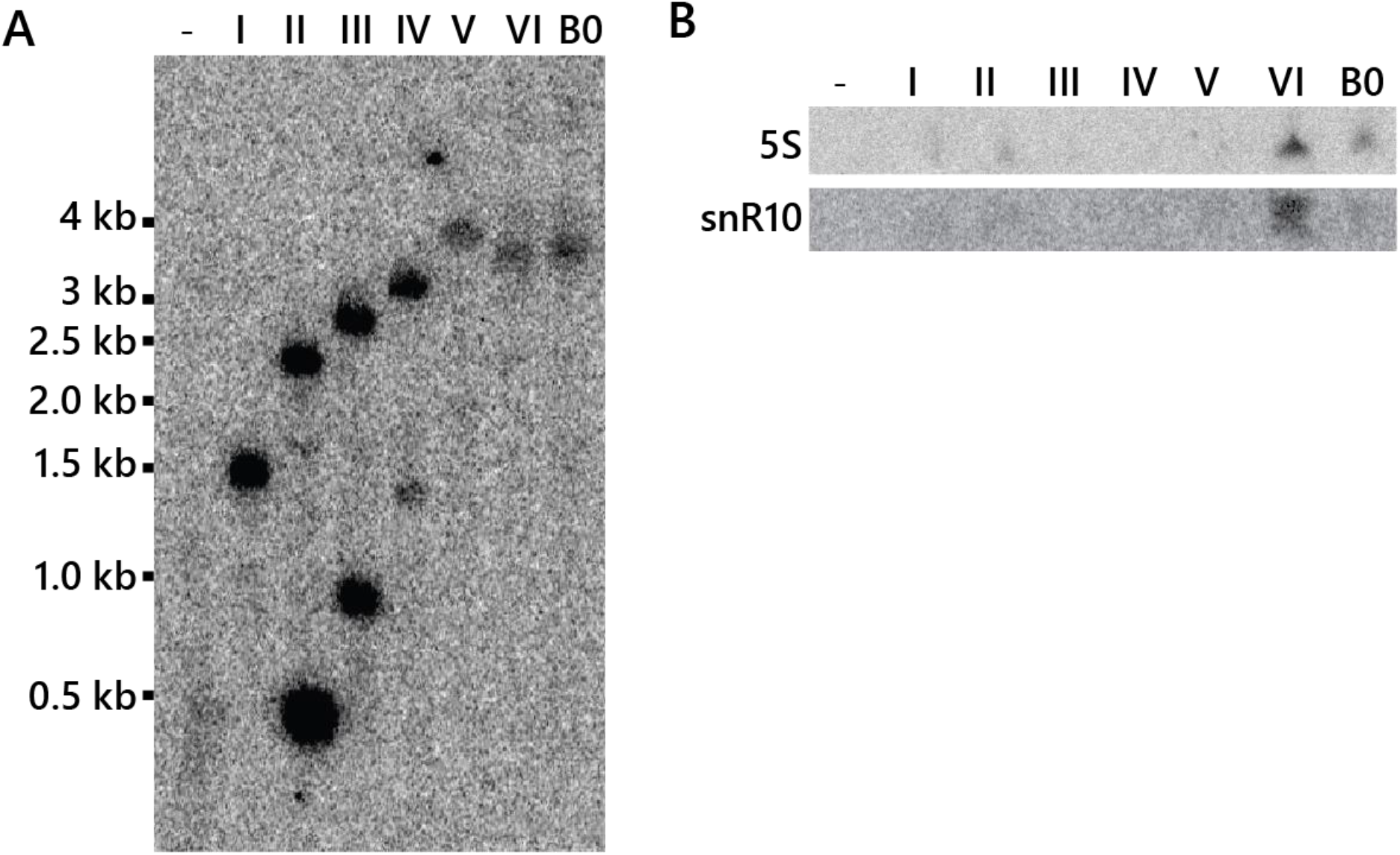
RNA analysis of pre-ribosomal complexes. (**A**) Northern blot with MS2 probe of total RNA extracted from cells where truncated pre-rRNA were expressed. (**B**) Northern blot with 5S and snR10 probes of RNA extracted from purified pre-ribosomal complexes.

**Supplemental Figure 2.**
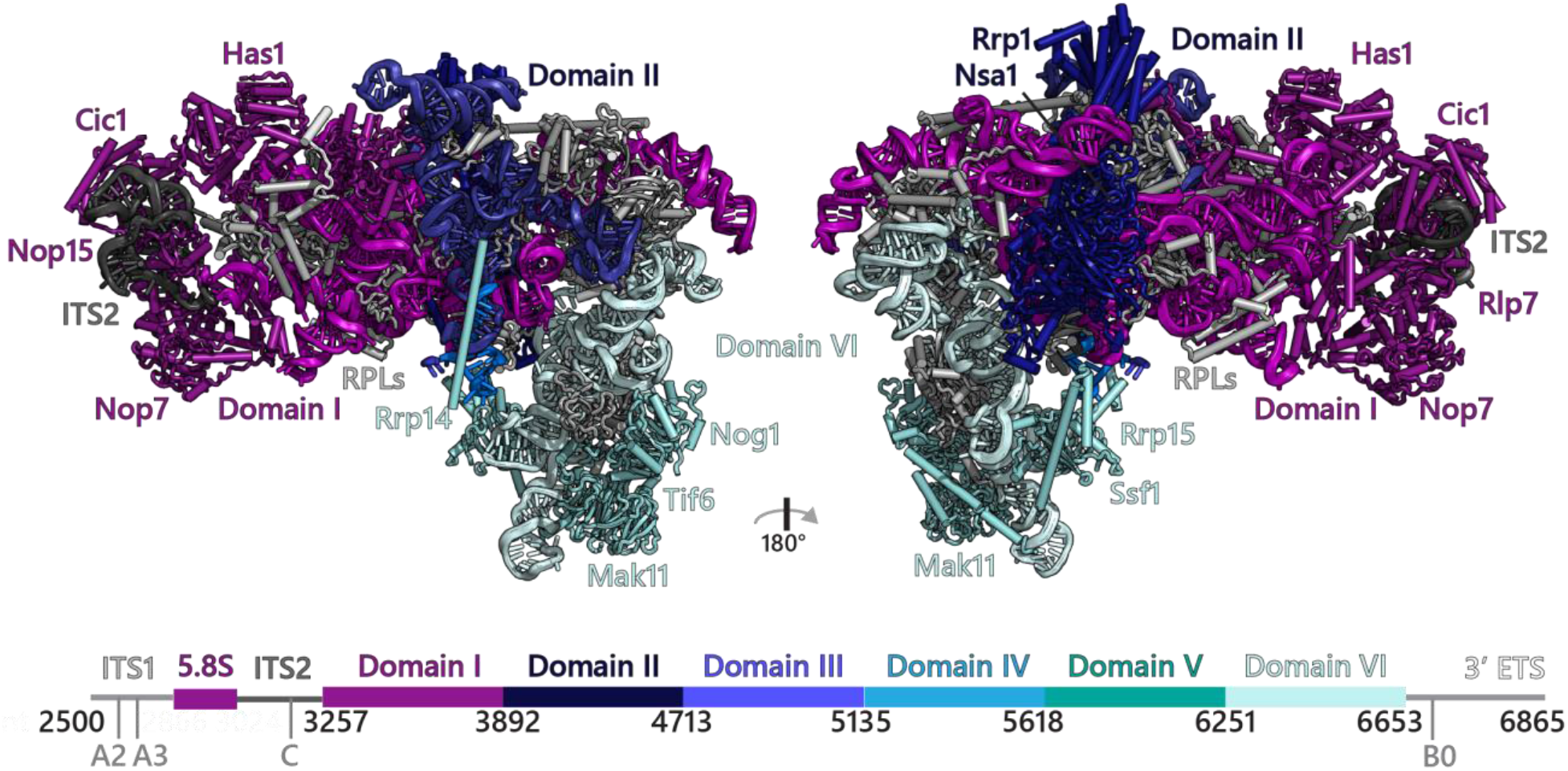
Mapping the recruitment of large subunit assembly factors on the nucleolar pre-60S particles. Model of state 2 of the nucleolar pre-60S particles (PDB 6C0F) (Sanghai et al. 2018) where the assembly factors are colored according to the stage to which they are recruited. Ribosomal proteins are shown in grey.

